# Lipid diffusion enhanced by membrane-associated molecular motors

**DOI:** 10.64898/2026.10.10.758165

**Authors:** Gwendal Guerin, Emily W. Sørensen, Ching-Yun Chen, John Manzi, Julien Berro, Nikos Hatzakis, Hsuan-Yi Chen, Feng-Ching Tsai

**Affiliations:** Laboratoire Physico-Chimi Curie, Institut Curie, Center de Recherche, Paris Sciences et Lettres Research University, Centre National de la Recherche Scientifique, 75005 Paris, France; Department of Chemistry and Nano-Science Center, University of Copenhagen, National Central University, Taoyuan City, 32001, Taiwan; Department of Physics and Center for Complex Systems, National Central University, Taoyuan City, 32001, Taiwan; Department of Molecular Biophysics and Biochemistry, Yale University, New Haven, CT, USA; Nano biology Institute, Yale University, West Haven, CT 06516, USA; Department of Cell Biology, Yale University School of Medicine, New Haven, CT 06520, USA; Institute of Physics, Academia Sinica, Taipei City, 11529, Taiwan

## Abstract

We experimentally investigate fluid lipid membranes coupled to nematic actin filament layers through myosin I motors. The motor activity of myosin I re-organizes the actin layer, modifying the morphology of +1/2 topological defects and generating spatially heterogeneous, actin-rich regions. Single-particle tracking (SPT) of lipid molecules reveal two interesting features. First, by identifying two lipid populations characterized by low- and high-speeds, it reveals an increased fraction of high-speed lipids in actin-rich regions. Second, by classifying short-time lipid trajectories into normal diffusion, directed motion, confined motion, and subdiffusion, it shows that lipids in actin-rich regions have a lower probability to stay in the confined and subdiffusion states, thereby favoring normal diffusion. Consistently, fluorescence recovery after photobleaching (FRAP) measurements show that lipid diffusion in actin-rich regions is enhanced. Together, these results suggest that myosin I motor activities redistribute lipids toward faster-moving population, and less confined and subdiffusive motion. As a result, lipid diffusion is enhanced by motor activity.

## Introduction

It has been known for almost three decades that when bacteria E. coli were added to a freely standing soap film, the two-dimensional diffusion constant of passive tracers in the film was significantly enhanced due to the flow produced by the swimming bacteria [1]. It has been known for almost three decades that when bacteria E. coli were added to a freely standing soap film, the two-dimensional diffusion constant of passive tracers in the film was significantly enhanced due to the flow produced by the swimming bacteria [1]. In principle, the same physical mechanism could be detectable at the molecular scale. For example, the activities of molecular motors in a two-dimensional fluid membrane should be able to produce small-scale random flows that enhance the diffusion constant of the lipids or other macromolecules in the membrane [2]. However, so far, such motor activity-enhanced diffusion in membranes has not been observed experimentally. Although there are high-precision single-particle tracking data on membrane proteins in living cells [3], the complexity of the cell membrane makes it difficult to reveal how motor activities affect lipid diffusion. Experimental studies on artificial membranes with a few components are more suitable for this purpose.

Ion pumps embedded in membranes are not the best candidate to produce diffusion-enhancing membrane flow because they exert forces mainly in the direction perpendicular to the membrane plane [4]. Motors with transmembrane domains that exert horizontal forces on membranes are more suitable [5], but such machines are challenging to reconstitute in well-defined model systems [6][7]. Studies using reconstituted systems have shown that when a thin layer of polymers, such as microtubules or actin filaments, are anchored to membranes through membrane-bound stepping motors, the motors can generate forces parallel to the membrane surfaces, resulting in polymer gliding [8] [9] [10] [11]. While significant re-organization of the polymers by motor activity was observed, how the motor activity impacts lipid diffusion in these systems has not been explored.

A motor protein with a motor domain binds to a polymer and with a tail domain that binds to a lipid should be able to generate lateral force on the membrane, thereby affecting lipid diffusion. For example, type I myosin (myosin-I) motors link actin filaments to PI(4,5)P_2_ (PIP_2_) lipids in the membrane with high affinity [12] [13] [14]. The motor domain of a myosin-I cyclically hydrolyzes adenosine triphosphate (ATP) when binding to both actin and membrane [15] [16]. The mechanical lever arm of myosin-I links the motor and tail domains to amplify its mechanical motion during ATP hydrolysis. Thus, unlike myosin II motors that are embedded within the cortical actin networks, myosin-I motors are located at the actin-membrane interface and generate forces on both lipids and actin filaments along the director of the polar filaments. Reconstituted systems with lipid membranes, actin filaments, and myosin-I motors are thus suitable for investigating whether and how the activity of membrane-associated motors impact lipid diffusion.

Myosin-I motors play key roles in various cellular processes involving dynamic reorganization of both actin and membranes, including cell migration, phagocytosis, and intracellular trafficking [12].Studies using reconstituted systems have revealed the ATP hydrolysis cycles of myosin-I, and its sensitivity to external force loads [17]. Myosin-I has been shown to reorganize branched actin networks [19][20] and accelerate filament depolymerization [21]. On fluid membranes, myosin-I can apply forces on the order of 1 pN on actin filaments [18]. Multiple myosin-I motors attached to the surface of giant unilamellar vesicles can generate 10 pN of forces to deform vesicle membranes into tubes by propelling along polar actin bundles [22]. Despite these insightful findings, the mechanical impacts of myosin-I motors on membrane dynamics remain unclear.

In this letter, we performed an experimental study of a supported lipid membrane coupled to a dense nematic actin layer through myosin-I motor, myo1b (Fig. 1a). We find that active myo1b motors reorganize the actin layer, generating lane-like and protrusion-like structures enriched in actin (Fig. 1c). These myo1b-driven actinrich structures are indications of local motor activity. Single lipid tracking (SPT) of DOPE lipids reveal two lipid populations with high and low mean speeds. Comparing regions of high and low actin density, we found that the fraction of high-speed lipids is higher in actinrich regions. Further analysis of lipid trajectories reveal lipids switch among diffusive, directed, confined and sub-diffusive states, and lipids in actin-rich regions exhibit a lower probability of being in the slow-diffusing confined and subdiffusive states. Consistent with the SPT measurements, fluorescence recovery after photobleaching (FRAP) measurements show that the effective diffusion coefficient of PIP_2_ lipids is higher in actin-rich regions than in regions with low actin density. Taken together, our results demonstrate that myosin-I motor activity at the actin-membrane interface re-organizes actin while enhancing local lipid diffusion.

**FIG 1.**
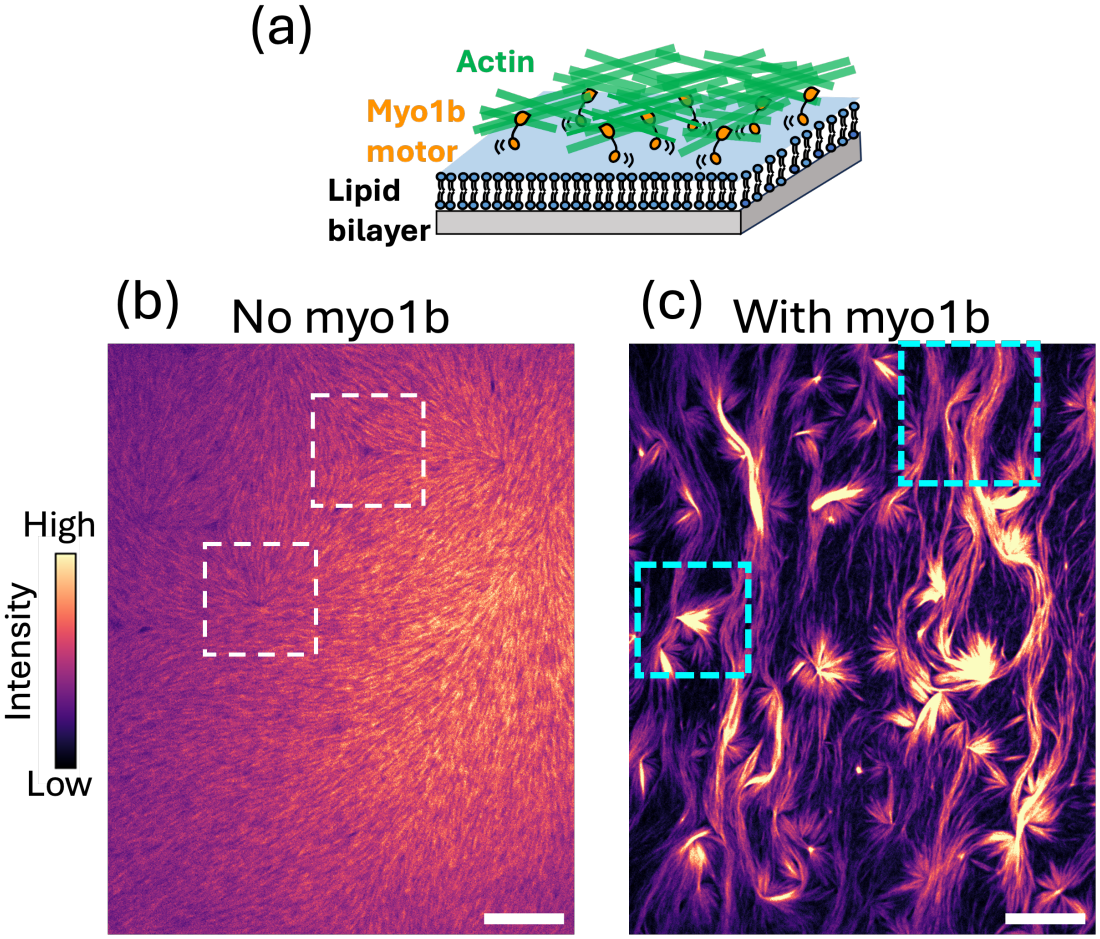
Myo1b motors reorganize an actin nematic layer on a planar membrane. (a) Schematic of the experimental setup: an actin layer coupled to a supported lipid bilayer (SLB) through myo1b motors. (b) Representative confocal fluorescent images showing an actin nematic on SLB in the absence of myo1b. White rectangles indicate the -1/2 (Top) and +1/2 (Bottom) topological defects. (c) Myo1b drives actin reorganization into lane-like (top box) and protrusion-like (bottom box) structures. Scale bars: 20 *µ*m.

## Results

Glass-supported lipid bilayers (SLBs) containing PIP_2_ covered by a dense layer of actin filaments were prepared (Fig. 1b) [23]. The filaments were stabilized by fluorescently tagged phalloidin (AX647-phalloidin), enabling visualization by fluorescent microscopy. In the presence of depletion agent methylcellulose, the filaments self-organize into a two-dimensional nematic on the membrane, exhibiting comet-like +1/2 and trefoil -1/2 topological defects (Fig. 1b). In the presence of myo1b at the membrane-actin interface, we observed morphological transformation of the actin nematic, exhibiting the accumulation of actin into lane-like and protrusion-like structures (Fig. 1c).

To investigate whether the activities of myo1b can impact lipid dynamics, SLBs supplemented with a small fraction of Atto647N DOPE, which do not bind to myo1b, were prepared. The trajectories of individual DOPE lipids were recorded with a frame rate of 100 Hz using Total Internal Reflection Fluorescence (TIRF) mi-croscopy [24]. Actin intensity thresholds 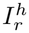and 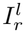 were used in order to compare the dynamics of single lipids in high actin intensity (intensity 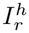; ”on actin”) and low actin intensity (intensity 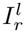 ; ”off actin”) regions. Following this actin intensity threshold, lipid trajectories were first classified into ”on actin” and ”off actin”. Then, we calculated the mean speed of the lipids along each trajectory, 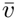, where *v* is the speed calculated from the displacement of two consecutive frames in the trajectories. In the following, data for systems with myo1b motors activity are presented to compare the behaviors of the lipids in actin-rich region, ”on actin”, and in regions with lower actin intensity, ”off actin”.

As shown in Fig. 2, the distribution of lipid mean speeds shows two populations, high-speed and low-speed, separated by a threshold speed *∼*15*µ*m*/*sec. We found that in actin-rich regions, the fractions of lipids in the high-speed population is higher (Fig. 2).

**FIG 2.**
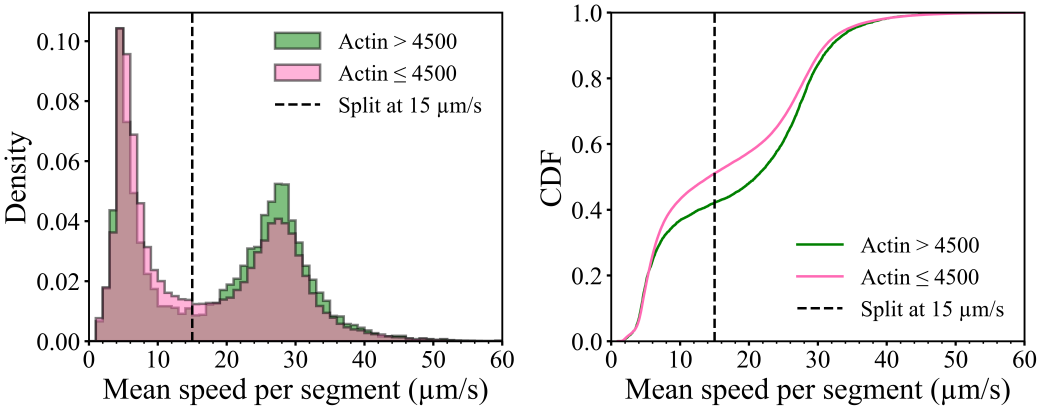
Left: Single lipid tracking reveals distributions of the mean lipid speed in the actin-rich region, ”on actin” (5569 tracks), and regions with lower actin density, ”off actin” (90230 tracks). Actin threshold intensity 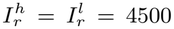. Right: Cumulative distribution functions (CDF) reveal differences in the probability of lipids belonging to the high-speed and low-speed populations for ”on actin” and ”off actin” lipids.

To determine how individual lipid motion differs between ”on actin” and ”off actin” regions, we analyzed single lipid trajectories using our recently developed machine learning algorithm DeepSPT [25]. DeepSPT classifies segments of a lipid trajectory into four moving states: normal diffusion (state 1), directed motion (state 2), confined (state 3), and subdiffusive motion (state 4). We defined the mean instantaneous speed 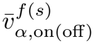 as the average instantaneous speed of fast (slow) lipids in state *α* (*α* = 1, 2, 3, or 4), in on-(off-) actin regions, with 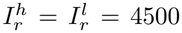. As shown in Fig. 3, 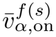is approxi-mately equal to 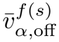 for a given moving state and speed population. These results indicate that myo1b activity does not alter the dynamics of lipids in a given fast/slow population and moving state.

**FIG 3.**
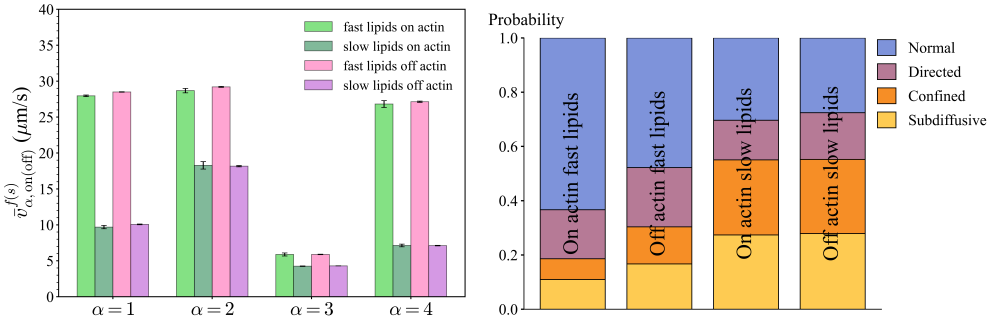
Left: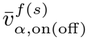, mean instantaneous speeds of lipids in each moving state. Right: Probabilities of lipid moving segments in a given fast/slow population and moving state.

We found that myo1b activities has a pronounced effect on the distribution of lipids among the moving states. As shown in Fig. 3, the fast-moving lipids in actin-rich regions has a higher probability to be in the normal diffusion state , with correspondingly lower probabilities of subdiffusive and confined states.

Together, the SPT data provide a coherent physical picture across a broad range of spatial and temporal scales. Myo1b motor activity redistributes lipids away from confined and subdiffusive states toward normal diffusion state, and increases the fraction of lipids in the high-speed population. We therefore expect this motor-induced enhancement of lipid mobility to extend from the single molecule level to collective lipid diffusion.

To probe lipid dynamics at the ensemble level, we performed FRAP measurements of fluorescently labelled PIP_2_ (TF-PIP_2_), which binds to myo1b, in regions of high and low actin intensities. As shown in Fig. 4, TF-PIP_2_ diffuses on average 17% faster in actin-rich regions (Fig. 4, a and c, ”Active myo1b”). We next tested whether this enhanced diffusion originates from myo1b motor activity by replacing ATP with the non-hydrolyzable analog AMP-PNP. In the absence of ATP hydrolysis, myo1b induced only local actin accumulation but not the large-scale protruding structures observed with ATP (Fig. 4d). In contrast to the ATP condition, FRAP measurements with AMP-PNP condition showed no significant difference in TF-PIP_2_ diffusion between regions of high and low actin intensity (Fig.4a, ”Non-active myo1b”). Together, these results show that myo1b motor activity drives actin reorganization while locally enhances lipid diffusion, consistent with the microscopicpicture obtained from single lipid analysis.

**FIG 4.**
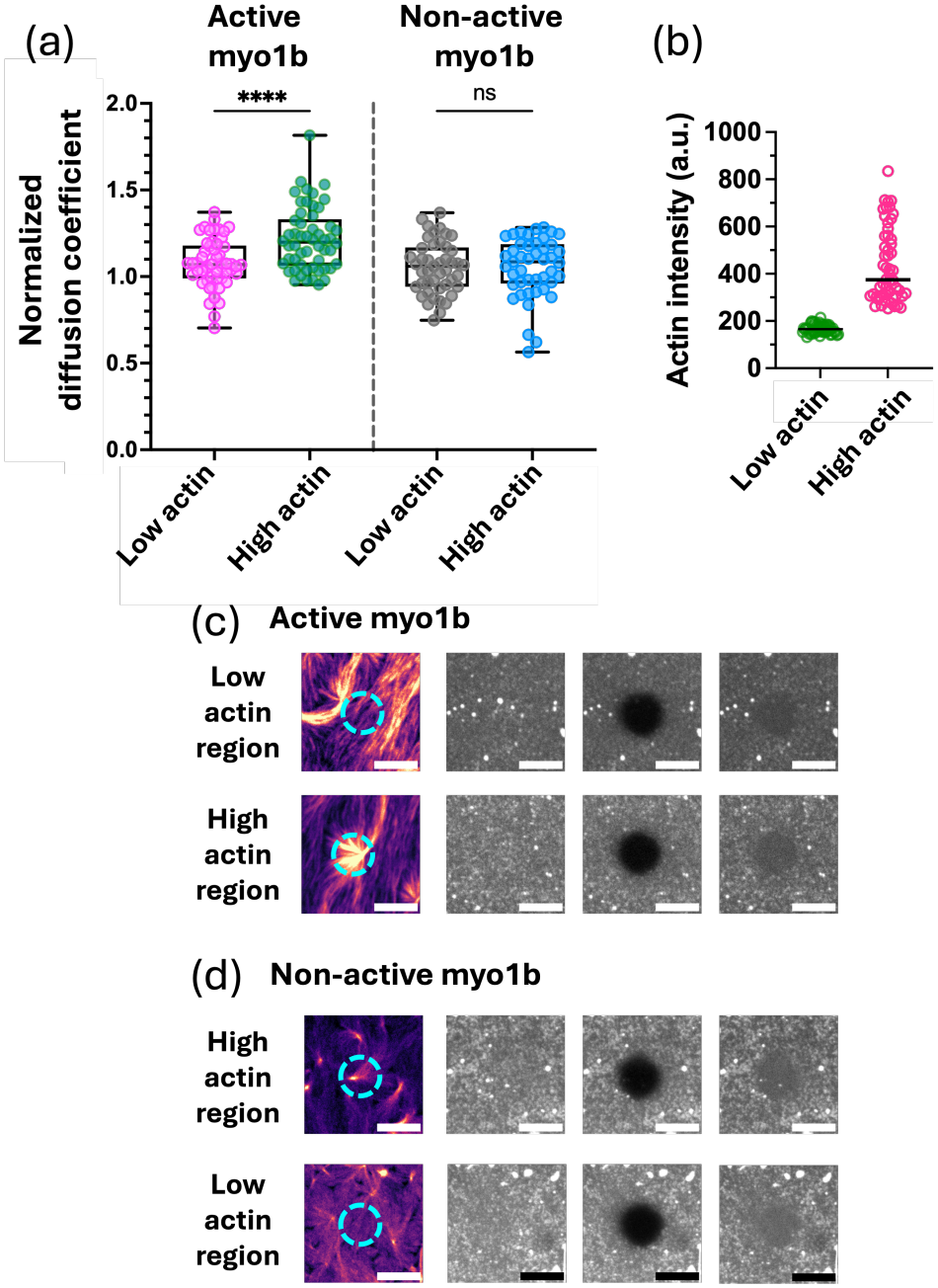
Myo1b motor activity enhances lipid diffusion in actin-rich regions. (a) Normalized diffusion coefficients of TF-PIP_2_ measured in low and high actin intensity regions for active and non-active myo1b conditions. Active myo1b: *N* = 4 independent experiments, and *n* = 54 FRAP measurements for both low and high actin regions. Non-active myo1b: *N* = 3 independent experiments, and *n* = 45 FRAP measurements for both low and high actin regions. For each experiment, diffusion coefficients were normalized with the median value of diffusion coefficients obtained from a bilayer in the absence of proteins. Statistic test: one-way ANOVA,****:*p <* 0.0001, ns: *p >* 0.9999. (b) Actin intensity in high and low actin regions, in active myo1b condition. *n* = 54 and 54 data points. (c and d) Representative confocal images of actin and FRAP measurements in Active myo1b (c) and Non-active myo1b (d) conditions. Scale bars: 10 *µ*m. Cyan circles indicate FRAP regions within a 10 *µ*m-diameter circular regions.

## Discussion

Our results reveal motor-driven lipid dynamics in a heterogeneous membrane environment. Individual lipids switch among different moving states along their trajectories, and the distribution of trajectory-averaged speeds exhibit two distinct populations. These observations suggest that the membranes contain dynamically distinct regions associated with fast and slow lipid motion.

The four moving states observed within individual lipid trajectories suggest that lipids move in a heterogeneous and fluctuating environment. In this picture, transient trapping can give rise to confined and subdiffusive motion, whereas random thermal force and local active perturbations contribute to diffusive and directed motion. This coexistence of dynamic states is similar to what were observed in dense colloids, dusty plasma, and supercooled liquids [26].

In summary, our experiments demonstrated the effect of myo1b motor activity on lipid motion from the molecular level to collective dynamics. Previous studies have shown that actin networks coupled to membranes through passive actin-membrane linkers slow down lipid diffusion [27]. Our findings suggest that active myosin-I-mediated actin-membrane coupling may provide cells a way to modulate local lipid dynamics.

The physical picture revealed in this study in which motor activity enhances lipid diffusion in a heterogeneous membrane is different from the theoretical model of mo-tors stirring a homogeneous membrane [2]. Instead, our results suggest that individual lipids explore a heterogeneous, time-dependent environment, and myo1b motor activity affects this environment by changing the occupancy of different moving states and the relative populations of fast and slow lipids. A more detailed quantitative analysis of the single-particle trajectories should be able to make a connection between this complex potential landscape experienced by the lipids and the effect of motor activity induced lipid flow. Such future works will help us to better understand active membrane dynamics at the kinetic level.

## Acknowledgment–

We thank Aurelie Bertin (i2BC, Paris Saclay) for her help with the filament length analysis and Frederic Pincet (ENS, Paris) for his help with the FRAP analysis. We thank Patricia Bassereau (Curie Institute, Paris) for fruitful discussions. F.-C.T. is a member of the CNRS consortium Approches Quantitatives du Vivant (AQV), Labex Cell(n)Scale (ANR-11-889 LABX0038) and Paris Sciences et Lettres (ANR-10-IDEX-0001-02). G.G. and F.-C.T. are supported by ANR grant ANR-20-CE11-0010-01 (ActinFission) and the Fondation pour la Recherche Médicale (FRM) (FRM EQU202003010307). G.G. and F.-C.T. and J.B. are funded by a PSL-Qlife grant (MyoMemActin). F.-C.T. is funded by ANR grant ANR 25-CE30-4672-01 (Active-ActinMembraneFlows). The authors greatly acknowledge the Cell and Tissue Imaging (PICT-IBiSA), Institut Curie, member of the French National Research Infractucture France-BioImaging (ANR10-INBS-04).

## Notes

### Competing Interest Statement

The authors have declared no competing interest.

### Summary of Updates

Manuscript text is updated for correcting typo and for clarity

